# Filming space-time changes of gene expression with *expressyouRcell*

**DOI:** 10.1101/2022.08.04.502810

**Authors:** Martina Paganin, Toma Tebaldi, Fabio Lauria, Gabriella Viero

## Abstract

The last decade has witnessed massive advancements in high-throughput techniques capable of producing quantifications of transcript and protein levels across time and space, and at high resolution. Yet, the large volume of big data available and the complexity of experimental designs hamper an easy understanding and effective communication of the results.

Here we present *expressyouRcell*, a unique and easy-to-use R package to map the multi-dimensional variations of transcript and protein levels in cell-pictographs. These variations are outcomes of differential and gene set enrichment analysis across space and time. Our tool directly associates these results with up to twenty specific cellular compartments, visualising them as pictographic representations of four different cellular thematic maps. *expressyouRcell* visually reduces the complexity of displaying gene expression and protein level changes across multiple time-points by generating dynamic representations of cellular pictographs.

We applied *expressyouRcell* to six datasets, demonstrating its flexibility and usability in the visualization of simple and highly effective static and dynamic representations of time course variations in gene expression. Our approach complements classical plot-based methods for visualization and exploitation of biological data, improving the standard quantitative interpretation and communication of relevant results.

## INTRODUCTION

In the world of big data we are living in today, visualisation tools and technologies are essential to analyse massive amounts of information, guide data-driven decisions and allow deeper understanding of the complexity of biological systems in physiological and diseased conditions^1–4^. During the last two decades, the generation of sequencing-based big data has witnessed an unprecedented explosion and a massive increase in the amount of raw information at our disposal^5–10^. As the complexity in the experimental designs - characterised by multiple variables, factors or covariates (e.g., conditions, time points and tissues) - increases, the biological data mining process has become more challenging. Thus, the scientific community requires next-generation visualisation tools and approaches to support the effective interpretation and intuitive presentation of results from complex experimental designs.

Typically, the biological meaning of sequencing data is assessed through differential analyses and downstream pipelines, such as annotation enrichment analysis, coupled with network and clustering analysis. However, with complex experimental designs, these approaches are of difficult interpretation, and they do not always satisfactorily disentangle the biological meaning of the data nor can effectively and rapidly communicate biological results.

Humans are visual organisms, as Aristotle said in his Metaphysics: “All men naturally desire knowledge. An indication of this is our esteem for the senses; for apart from their use we esteem them for their own sake, and most of all the sense of sight.” (Aristotle, Metaphysics ∼ 400 B:C). Today more than ever, the visualisation of complex data in simple and information-rich graphical formats are of the utmost relevance to visually exploit complex abstract information. The overarching aim is to fulfil the double ambitious purpose of quantitative data mining and effective communication in a blink of an eye.

Dedicated graphical tools have been proposed as means to represent biological information through graphical approaches in the past^11–15^. Available web applications, such as GeneCards, UniProt or Human Protein Atlas^11,14,15^ mainly consist of a user-friendly front-end for exploring gene and protein expression data obtained from experiments performed in various tissues, cell types and species. To highlight the localization of genes/proteins, the experimental results can be visualised using schematic representations of cells or organisms^11,14,15^. However, these tools lack versatility. In fact, they allow the users to just browse the information available in the database rather than customise the visualisation of his/her own data. An R package and accompanying web application have been developed to draw representations of expression datasets in human and mouse tissues^13^. However, this tool requires a series of advanced computational steps to handle and visualize the user’s own data^13^. Recently, an R-based application has been proposed to compute and represent results from enrichment analyses through pictographs^12^, but none of these tools reduce the biological complexity of results from sophisticated experimental designs. The lack of efficient tools for next-generation data exploitation is an urgent need in biology today.

To portray quantitative gene expression changes in space and time in a very intuitive and effective manner, we propose *expressyouRcell*, an user-friendly and flexible R package that generates both static and dynamic cellular pictographs using custom-made complex NGS experiments. Leveraging the concept of choropleth maps, we used specific types of thematic maps for representing variations of variables, i.e., fluctuations in gene and transcripts expression levels. These values can be directly mapped to specific cellular compartments and visualised in pictographic representations of multiple cell types. *ExpressyouRcell* also generates movies of dynamic changes in the cellular pictographs at subcellular and organelle-level resolution. This function is uniquely suitable for visualising fluctuations in gene expression levels across multiple time points or to represent outcomes from differential longitudinal analyses. *expressyouRcell* unlocks an enormous simplification of data understanding and a conceptual shift as to how scientific data communication of biological results can be achieved.

## MATERIALS AND METHODS

### User’s input data

*expressyouRcell* is optimised for representing multiple gene expression datasets and accepts lists of tables. Each data table must contain a set of gene symbols which is sufficient to perform gene set enrichment analysis and visualise these results through cellular pictographs.

Additionally, the gene symbols can be coupled with: i) their expression level expressed in read counts, count per million of reads (CPM) or reads per kilobase of gene per million (RPKM) or ii) fold-changes (FC) and p-values from upstream differential analyses. FC are particularly useful to highlight the most affected cellular compartments and structures across different experimental conditions, time points and tissues.

To improve the flexibility in handling different data structures, *expressyouRcell* provides the user with multiple options for colouring subcellular localizations (see details in section *Visualizing gene expression data to the cellular pictograph* for details).

### Gene localization mapping

To define the colour and the shade of cellular pictographs, each gene must be previously mapped to a specific subcellular compartment. The localization of genes can be either provided by the user or assessed by *expressyouRcell* through the dedicated *map_gene_localization* function. This function requires as input a gene annotation file, provided in GTF format, and performs gene ontology enrichment analysis on the sets of input gene symbols. Single or multiple terms of the cellular component ontology, either cellular compartment or macromolecular complex, will be assigned to each gene.

The subcellular compartments drawn in the pictographs are described by terms from the cellular component ontology. Since the logical data structure of the gene ontology is organized as an hierarchical tree graph, we selected a subset of higher-ranked terms as descriptors for the pictographic organelles and compartments.

### Cellular pictograph drawing

*expressyouRcell* offers pictographs of four cell types: i) a typical animal cell, ii) a neuron, iii) a fibroblast and iv) a microglial cell. The desired cellular type has to be specified by the user in the *color_cell* function parameters.

The spatial coordinates of regions/organelles are extracted from multiple vector graphic files, one for each subcellular region or organelle, through the *rsvg_svg* function of the *rsvg* R package. The resulting cellular structures are then stored in RData objects, provided by *expressyouRcell*, and used by the function *geom_polygon* from the *ggplot2* package to generate the final pictographs.

A selection of subcellular regions and organelles have been chosen and drawn to create the cell pictographs (**Table S1**). The selection was based on the highest ranked nodes in the gene ontology graph and includes the main cellular regions and organelles. The subset of cellular component terms used for the cellular pictograph is cell-type specific.

### Visualising gene expression data to the cellular pictograph

The *color_cell* function provides two main approaches for assigning colours to the subcellular localizations and for visualising gene expression data pictographs.

The first option is based on the outcome of gene set enrichment analysis and on its representation.

This approach takes into consideration the identity and localization of genes, but not their expression level. Starting from the complete list of gene symbols, *expressyouRcell* performs the gene set enrichment analysis for each cellular component and assesses the statistical significance of the enrichment by the Fisher’s test and the colour shade of each compartment is defined by its associated p-value. An optional gene classification can be specified by the user through distinct categorical variables (e.g., “down-regulated” and “up-regulated”). In this case, a cellular pictograph for each class is generated following the same procedure described above.

The second option allows the visualisation of either gene expression levels or the result of previously performed differential analyses. This approach requires, for each gene, a numerical value reporting i) its gene or transcript expression level (reads count, TPM, CPM or RPKM) or protein levels, or ii) the outcome of differential analyses among multiple samples (e.g., fold changes, fold enrichments). The name of the column that defines the colour of the pictograph must be specified by the user. To provide further flexibility in data visualisation, *expressyouRcell* can generate cellular pictographs on the entire set of genes or on subsets defined by the user. In the first case the colour of each cellular compartment is defined by the mean (**Equation 1**), or median, of all gene-specific values associated with each localization

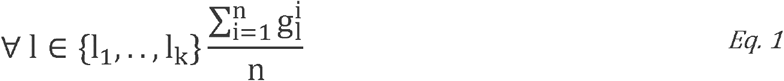

where *l* ∈ {*l*1, …, *lk*} is a cellular localization, *n* is the total number of genes associated with the localization *l* and g_i_ ^l^ is the gene-specific value (gene expression level or fold changes/enrichments). In the second case, e*xpressyouRcell* generates multiple pictographs, one for each subset of genes defined by the user according to an optional classification of genes (e.g., “down-regulated” and “up-regulated”). With this option, a cellular pictograph for each class is generated and the colour of each cellular compartment is calculated as in **Equation 2**, for each gene subset.

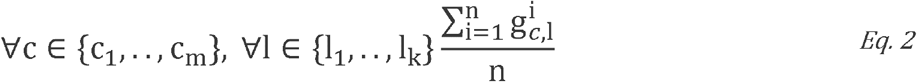

where *c* ∈ {*c*1,…, *ck*} defines a class of genes, *l* ∈ {*l*1, …, *lk*} is a cellular localization, *n* is the number of genes in class *c* associated with the localization *l*, and g^i^_c,l_ corresponds to their gene-specific value (gene expression level or fold changes/enrichments).

If no categorical variables are specified by the user, *expressyouRcell* classifies the genes according to a combination of cut-off values (e.g., *fold-change* and *p-value* thresholds).

### Animated cellular pictographs

*expressyouRcell* generates dynamic representations of data shown in the cellular pictographs. This is particularly useful when the user’s input data consists of multiple sets of time-course gene expression data. The generation of animated pictographs depends on information obtained by the *color_cell* function.

In this step, multiple data structures obtained by the *color_cell* function are required as input to the *animate* function, which creates a short movie or an animated picture. Additional mandatory inputs consist of the name of data tables for generating the animation, the transition duration (in seconds) and the number of frames/transitions. A bar timeline at the top of the animation plot allows a better tracking of the changes over time. Hence, the user can use as additional input a vector of labels for a customised timeline.

The working flow of this function begins with the generation of temporary frames with intermediate colour shades for each transition. The complete set of frame pictures is merged into a single animated picture (GIF) or short movie in *mp4* format. The *gifski* and *av* packages are used to produce the animated GIF picture and the movie, respectively.

### Output

*expressyouRcell* generates both static and dynamic representations of cellular pictographs that are associated with two main types of data structures.

The *color_cell* function generates static cellular pictographs and returns a list with multiple data structures. The first is a data table with six columns, reporting, for each subcellular component: i) its name, ii) the numeric value computed during the colour assignment step, iii) a numeric identifier for grouping the cellular localizations by colour, iv) its associated colour shade, the v) the identifier of each dataset, and, if present, and vi) the variable used for grouping the genes by classification.

The second data structure is a data table summarising the information on ranges used to categorise each subcellular localization (e.g., start, end, colour, and labels).

The third and fourth data structures are lists of graphical objects of class *ggplot*, and datasets used to plot the resulting cellular pictographs, respectively.

The *animate* function generates dynamic cellular pictographs and saves them either as movie (in *mp4* format) or as animated pictures (in *gif* format). The saving step is performed directly within the function.

## RESULTS

### Workflow

Here, we present *expressyouRcell*, an R package that generates static and dynamic cellular pictographs to represent outcomes of expression data analysis (RNAseq and proteomics) at sub-cellular and organelle-level resolution (**Figure 1A**). The input of *expressyouRcell* consists of a list of one or multiple tabular data structures with the biological information that the user desires to represent as cellular pictographs.

**Figure 1.**
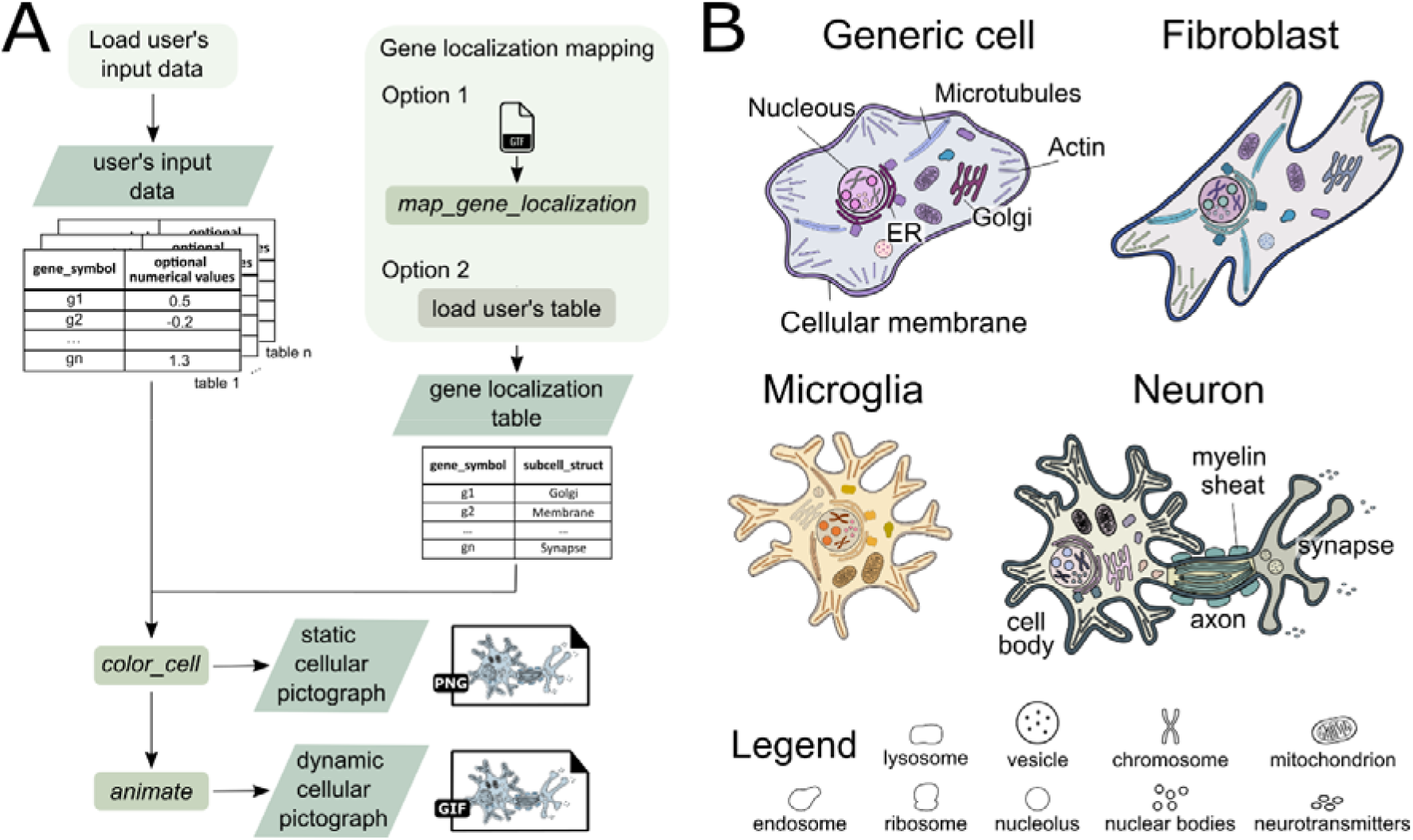
Workflow overview and cellular pictographs. (A) Flowchart representing the basic steps of *expressyouRcell*, the input requirements and the outputs. User’s input data are loaded within the package through the *color_cell* function, which also needs the *gene-localization* table. This data structure can be either generated through the *map_gene_localization* function (option 1) or provided by the user (option 2). The *color_cell* function assigns colours to cellular compartments according to i) the statistical significance of enrichment analysis, ii) fold changes from differential analyses or iii) gene expression or protein level abundances, averaged for genes in each compartment. Then, it outputs static cellular pictographs (in PNG format) and the data structure required to the *animate* function for the generation of animated cellular pictographs (in GIF or MPG format). User’s input data are defined in light green boxes. Diamond boxes denote intermediate and final output data. Functions provided within the package are indicated in italic font. (B) Set of available cellular pictographs, based on different cellular types: i) a generic animal cell, ii) a fibroblast, iii) a microglial cell and iv) a neuron. The chosen cellular pictograph has to be provided by the user to a dedicated parameter of the *color_cell* function. Default colours are assigned to organelles and subcellular compartments.

A table that associates genes ID with subcellular compartments is also required and can be either provided or generated through the *map_gene_localization* function that is available in *expressyouRcell*. The list of data tables and the gene localization table represent the input of the *color_cell* function, which assigns colours to cellular compartments according to the statistical significance of enrichment analysis, or based on fold change averages for genes in each compartment. Then, the function outputs static cellular pictographs and the data structure required to generate animated cellular pictographs or movies (through the *animate* function).

*expressyouRcell* provides the users with custom options for the generation of multiple static cellular pictographs or dynamic pictographs in the form of movies, and offers four thematic maps, based on different cellular types: i) a typical animal cell, ii) a neuron, iii) a fibroblast and iv) a microglial cell **(Figure 1B)**. Each cell type comprises eighteen organelles and macromolecular complexes (nucleus, Golgi apparatus, endoplasmic reticulum, cell membrane, cytosol, chromosome, vesicles, endosomes, cytoskeleton, lysosomes, mitochondria, neurotransmitters, ribosomes, nuclear bodies, nucleolus, extracellular matrix (**Figure 1B**). In addition, the neuron pictograph includes the myelin sheath and its global structure is organized into cell body, axon and synapse (**Figure 1B and Table S1**).

### Visualise longitudinal differential analysis

To demonstrate the flexibility and power of *expressyouRcell*, we present six case studies (**Table 1**), which represent particularly suitable applications of our tool. The selected studies share complex experimental designs with multiple variables, or covariates (e.g., conditions, time points and biological samples) based on two experimental techniques (RNA-Seq and mass spectrometry) and specific computational pipelines^5–10^.

**Table 1.** Summary of works and related datasets used to evaluate *expressyouRcell* features.

| Organism | Cells/tissues | # time points | Study title | Reference |
| --- | --- | --- | --- | --- |
| Mouse | Microglia | 4 | microglia activation in response to pathological tau | Wang et al., 2018 |
| Mouse | Neurons | 3 | RNA Expression in Excitatory Neurons during Hippocampal LTP | Chen et al., 2017 |
| Mouse | B/SC/M/H/K | 2 | Changes in GE during disease progression | Doktor et al., 2017 |
| Mouse | Schwann cells | 6 | Wound reprogramming and peripheral nerve regeneration | Clements et al., 2017 |
| Mouse | Neurons | 9 | temporal regulation of alternative splicing during neural development | Weyn-Vanhentenryck et al., 2018 |
| Human | hiPSCs | 7 | Protein levels in cells during differentiation from hiPSCs to hepatocyte-like cells | T. Hurrell et al., 2019 |

In the first case study, Chen and collaborators^6^ used TRAP-seq to characterise the transcriptome changes in a specific population of neurons after induction of long-term potentiation (LTP) at 30, 60 and 120 minutes after stimuli. The canonical analysis and data exploration with volcano plots displays the quantity and magnitude of differentially expressed genes (DEGs) identified between LTP and control condition at each time point, and the corresponding GO enrichment analysis (**Figure 2A and B**), demonstrating the complexity in data interpretation.

**Figure 2.**
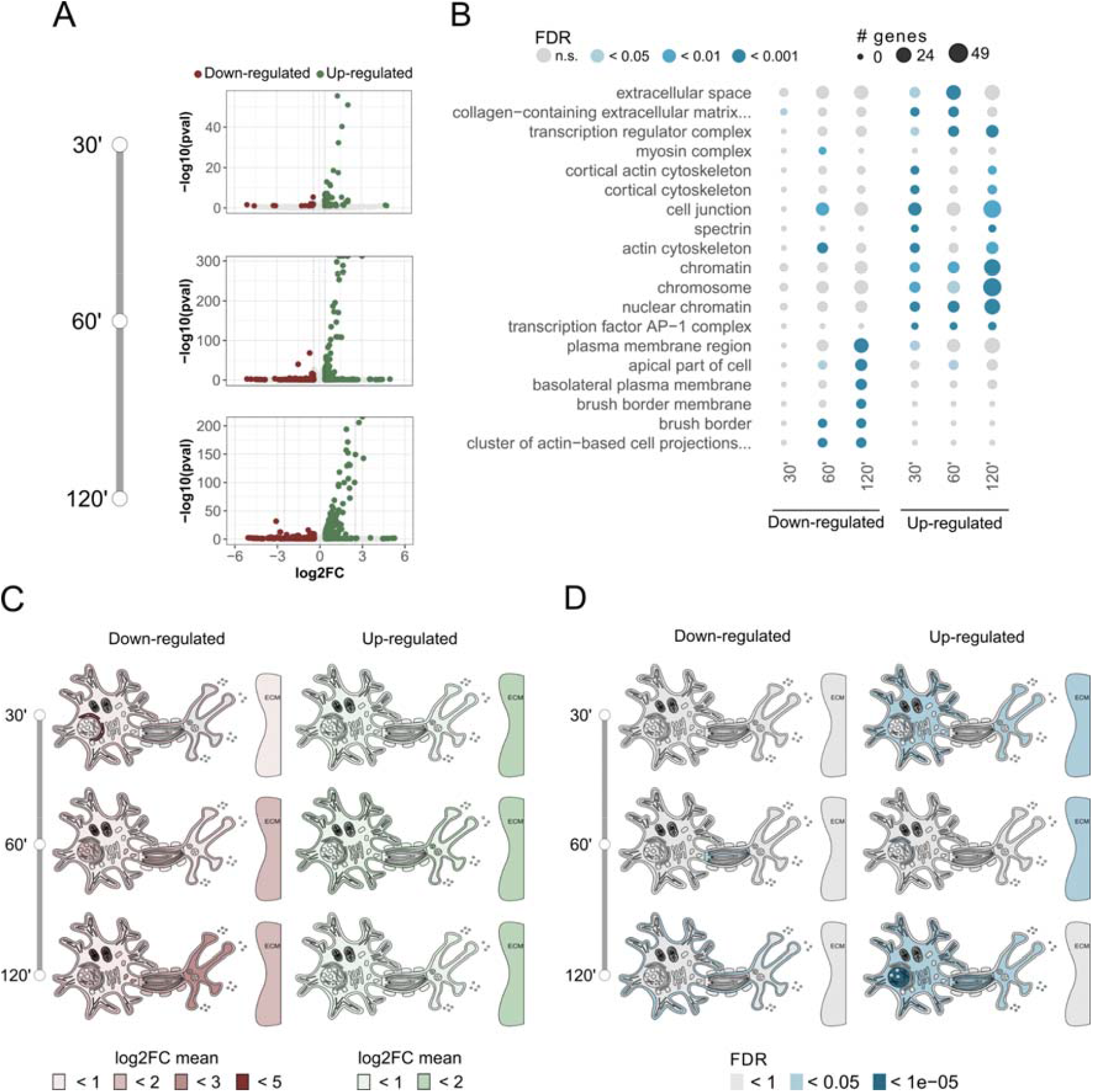
Standard approaches and cellular pictographs for visualising results of differential analyses across time-points. (A) Volcano plot of DEGs between LTP and control condition at indicated time point from TRAP-seq of neurons. Fold changes are plotted against the –log(p value). The vertical dashed lines indicate the 0.4 fold change cut-off thresholds. (B) Enrichment analysis of Gene Ontology (Cellular Component) terms at indicated time point of differentially expressed genes from TRAP-seq of neurons. Colours of dots indicate the significance of the enrichments. Dot sizes define the number of genes associated with each category. (C) Neuron pictographs from TRAP-seq. Colour shades of cellular compartments and organelles are based on logFC values from differential analysis. Up- and down-regulated genes were defined with the threshold described in (6) (i.e., abs(log2FC) > 0.4 and FDR < 0.1). (D) Neuron pictographs from TRAP-seq. Colour shades of cellular compartments and organelles are based on the significance of the enrichments. Up- and down-regulated genes were defined with the threshold described in (6) (i.e., abs(log2FC) > 0.4 and FDR < 0.1).

When we generated the neuron pictographs with *expressyouRcell* we clearly visualise at first glance: i) a gradual increase in the colour intensity over time, in particular for down-regulated genes (**Figure 2C**, red pictographs), and ii) changes in gene expression spreading across several cellular compartments (**Figure 2C and movies S1-S2**). With this visualisation we can appreciate the spatial localisation of gene expression changes and observe the emergence of down-regulated genes at 30’, with robust negative fold-changes encoding for proteins localised in the endoplasmic reticulum. At later time points, genes with negative fold changes are connected to mitochondria, nucleoplasm and the synaptic region of neurons.

We also visualised as heatmaps the results of gene set enrichment analyses on differentially expressed transcripts at all time points (**Figure 2B**). A significant enrichment of down-regulated genes for the cell membrane, which intensifies at the last time point (120’), can be swiftly observed. Up-regulated transcripts show a clear enrichment for chromatin and nuclear-related themes, in particular at 120 minutes after LTP induction. Next, we used *expressyouRcell* to display the results of gene set enrichment analysis through cellular pictographs of neurons for up- and down-regulated genes at each time point (**Figure 2D and movies S3-S4**). Coherently with the heatmap visualisation, the representation obtained with our tool shows very effectively and intuitively that down-regulated genes are enriched in the cell membrane theme at 120 minutes after LTP induction. From cellular pictographs generated for the enrichment analysis on up-regulated genes, we can observe that nucleoplasm and chromosomes are highlighted at all the time points, but at 120’ the enrichment is even more evident.

To appreciate the reduction in data complexity and the power of graphical representations provided by *expressyouRcell*, we analysed a second case study^5^. The dataset is characterised by higher complexity compared to the previous one in terms of number of time points. The experimental design consists of transcriptomes of mouse cortices at nine different time points during development^5^. It is clear that any pairwise comparison would generate three times more volcano plots and GO heatmaps than those shown in **Figure 2 A-B**. For each developmental time point we generated neuron pictographs (**Figure 3**) based on the expression level (expressed as transcript per million, TPM) of a subset of genes of interest selected by the authors^5^. We can very easily visualise and identify an increase in the TPM signal in myelin sheath-related genes starting from the fourth day after birth, accompanied by an increment of signal in the synaptic region. Applying the function movie, the results are even clearer and with strong communication impact (**Movie S5**).

**Figure 3.**
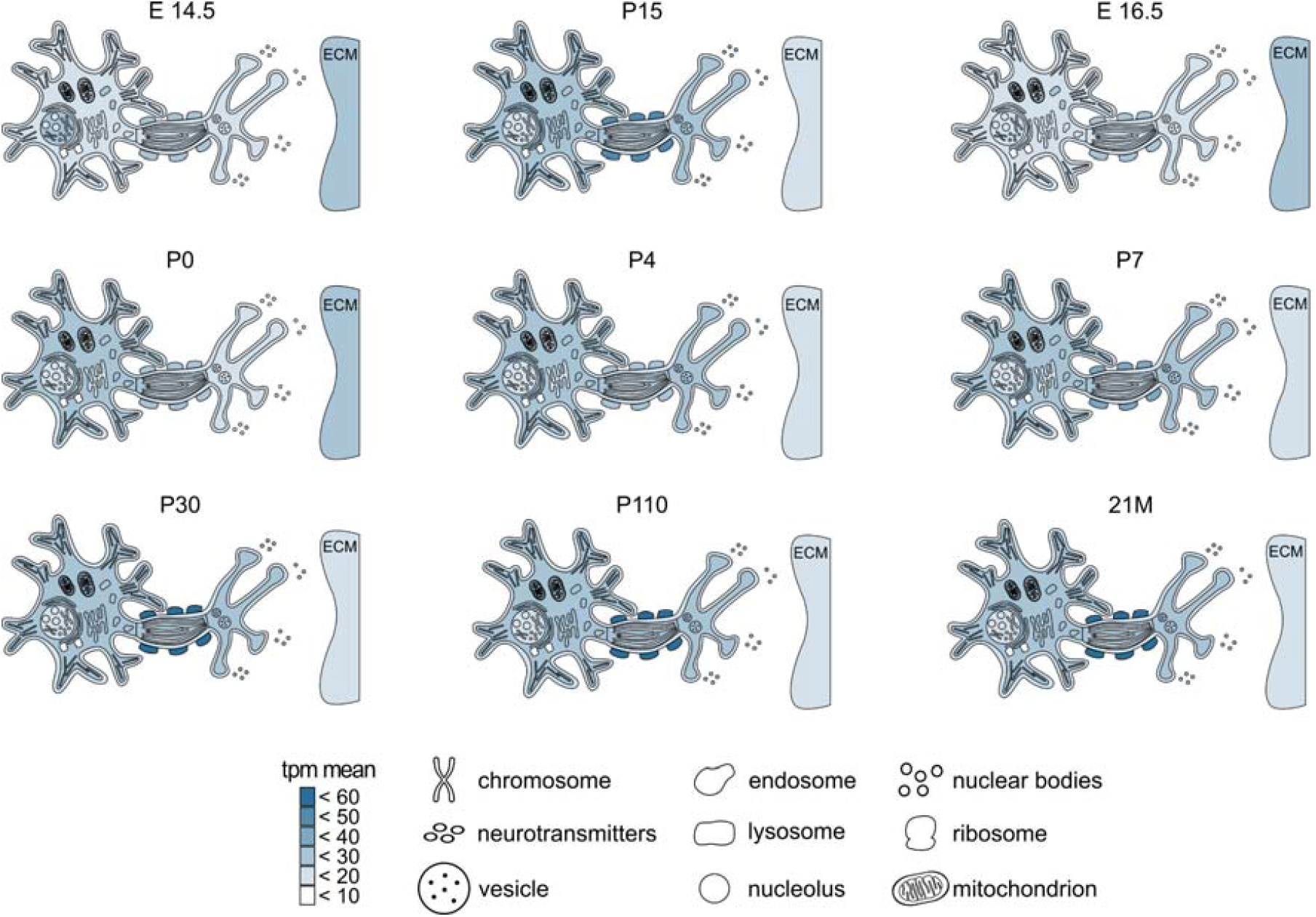
Neuron cell pictographs from RNA-seq of cortical neurons. Colour shades of cellular compartments and organelles are based on gene expression levels (measured in TPM), described in ^5^. Counts of RNA-seq data from this study were processed through https://www.refine.bio.

These results and those related to other case studies and applications of *expressyouRcell* (**Supplementary Figures S1-S7, and Movies S1-S17**) demonstrate that our tool is flexible and powerful allowing simple and intuitive data mining and visualisation for the next-generation of biological data exploitation.

## DISCUSSION

Differential gene expression outcomes (i.e., the absolute number of differentially expressed genes/transcripts, their magnitudes and significance values) are usually displayed through bar plots, volcano plots or heatmaps. As methodologies to peer inside the complexity of biological mechanisms evolve, the amount of biological information to be disentangled explodes and standard procedures are no more sufficient to fulfil this task. Hence, new visualization approaches become essential for supporting the interpretation of outcomes from data analysis pipelines and visually conveying meaningful information. For instance, volcano plots are used to display significant changes, but they are not suitable to represent the spatial localization of fluctuations in gene expression levels across cellular compartments. Hence, the canonical representation of differentially expressed genes (DEGs) through volcano plots can be integrated with intuitive and informative representations of the localization and intensity of changes provided by *expressyouRcell* pictographs.

Compared with standard methods, cellular pictographs generated through *expressyouRcell* offer an immediate approach to data exploitation, and a reduction of biological complexity. In particular, this method gives us the unique advantage of an immediate detection and intuitive illustration of the most affected cellular components, together with the intensity of the variations.

A handful of tools with alternative representation methods^11–15^ have already been proposed, but at the time of this writing, there are no comparable alternatives to *expressyouRcell*. In fact, our tool addresses the data visualization problem in a different and novel approach compared to existing ones^11–15^. It gives the unique possibility of creating dynamic illustrations, which can effectively support researchers in tracking the fluctuations in gene expression levels across multiple time points and reduces the complexity of biological information. Importantly, changes are dynamic by definition, and animated pictographs constitute a more appropriate solution for scientific communication than static illustrations. As of now, no other tool but *expressyouRcell* have been proposed to supply this need.

Some of the currently available tools mainly consist of web interfaces designed to query biological databases in a human interpretable manner, which comes particularly handy to retrieve information on single genes and proteins^11,14,15^. In particular, these web resources rely on pictographic illustration of cells to display the location of protein at subcellular resolution. As such, their purpose and functionality differ from our approach, while remaining useful resources with a wealth of biological information.

Two available R-based applications (i.e., *gganatogram* and *subcellulaRVis*) aim at representing biological data through cellular pictographs. *gganatogram* mainly produces anatomical pictographs for different organisms^13^. It offers nicely detailed pictographs of various whole-body organisms for visualizing biological data at organs and tissue level. The set of available pictographs also includes one generic cellular map complete with multiple organelles. Although the great advantage of visualizing biological data at the subcellular level, this option limits the users in representing specific cellular types with particular subcellular compartments. Furthermore, the tool requires a series of advanced computational steps to integrate and visualize the user’s own data. *subcellulaRVis* is a web R-based application which performs and visualizes results from enrichment analysis. This tool provides the users with two graphical options: a generic cellular pictograph and a more specific visualization of the endosomal system^12^ which is particularly useful to represent detailed and specific information for the endosomal compartments. The user-friendly web page interface offers the possibilities of performing enrichment analysis also to less-experienced users, and of generating easy-to-interpret graphical results. Unfortunately, this tool currently does not manage multiple data types, making it well-suited for performing only enrichment analysis.

Overall, *expressyouRcell* meets both users’ needs for dealing with a variety of different data types and their desire for customized visualizations. Our tool provides the scientific community with a fresh and original approach for representing gene expression changes and for effective, fast and dynamic communication of results through various cellular pictographs. This visualisation can support and complement the standard and already widely adopted methods with the additional knowledge on both the intensity and spatial localization of gene expression variations and open a completely new scenario as to how biological data exploitation can move towards the future.

## DATA AVAILABILITY

*expressyouRcell* is a package written in the R programming language which depends on multiple R packages, such as *ggplot2* for data visualisation, *clusterProfiler* for gene ontology enrichment analysis, *gifski* and *av* for the GIF animation and video realisation, respectively. *expressyouRcell* source code can be downloaded from the following GitHub repository: https://github.com/LabTranslationalArchitectomics/expressyouRcell, accompanied with installation instructions and functions documentation. *expressyouRcell* is an Open-Source software package (distributed under the MIT licence), which is compatible with Linux, Mac, or Windows PCs. Future releases may be expanded by additional cellular pictographs to support the visualisation of data on a wider range of cellular types. Also, an interactive web application could also be deployed through the Shiny framework to further expand the usability and reachability of our tool.

The package includes the R implementation of *expressyouRcell*, data used in this article, extensive documentation and a stable release.

## ACCESSION NUMBERS

This paper analyses existing, publicly available data. The accession numbers for the datasets are listed in the following table.

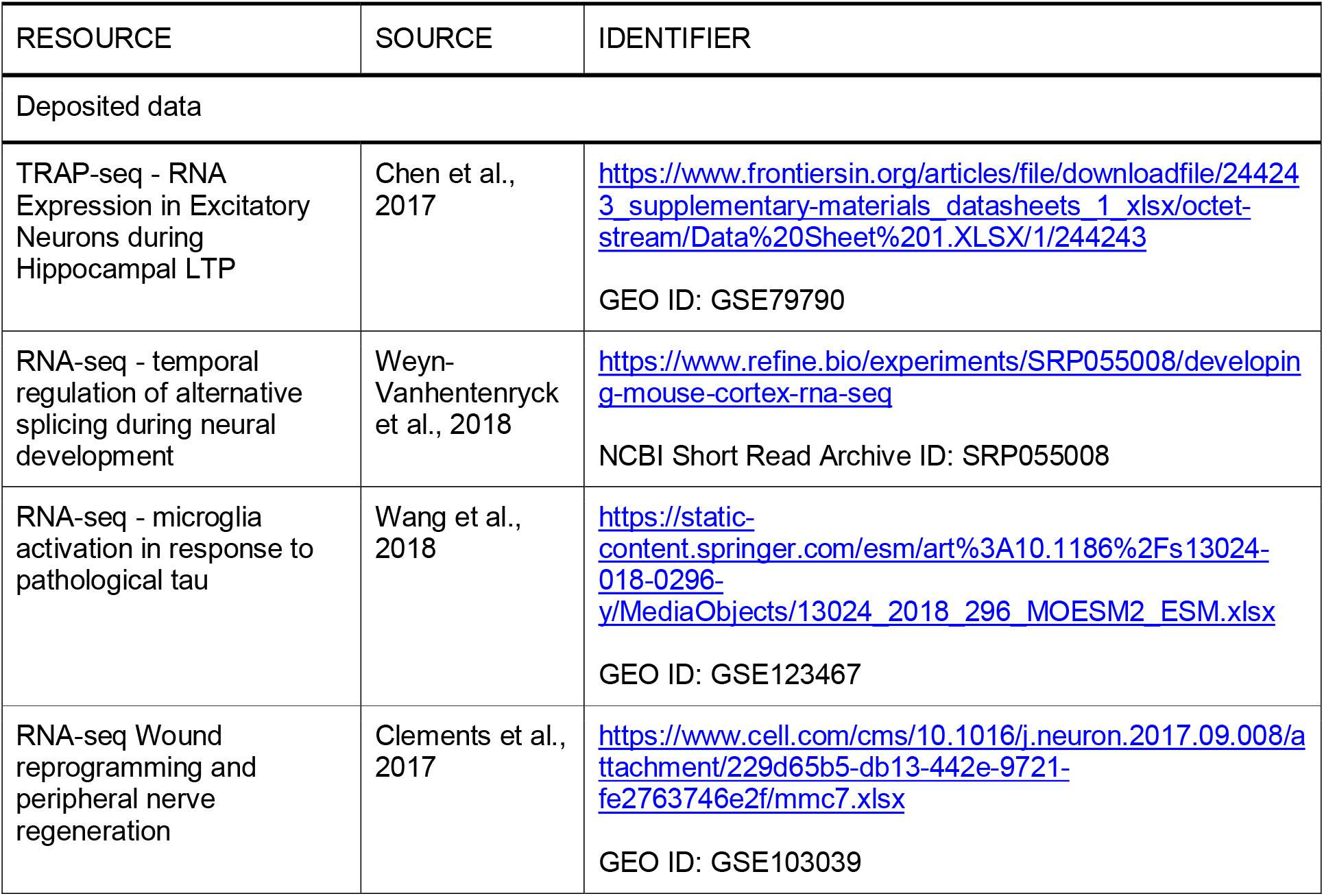
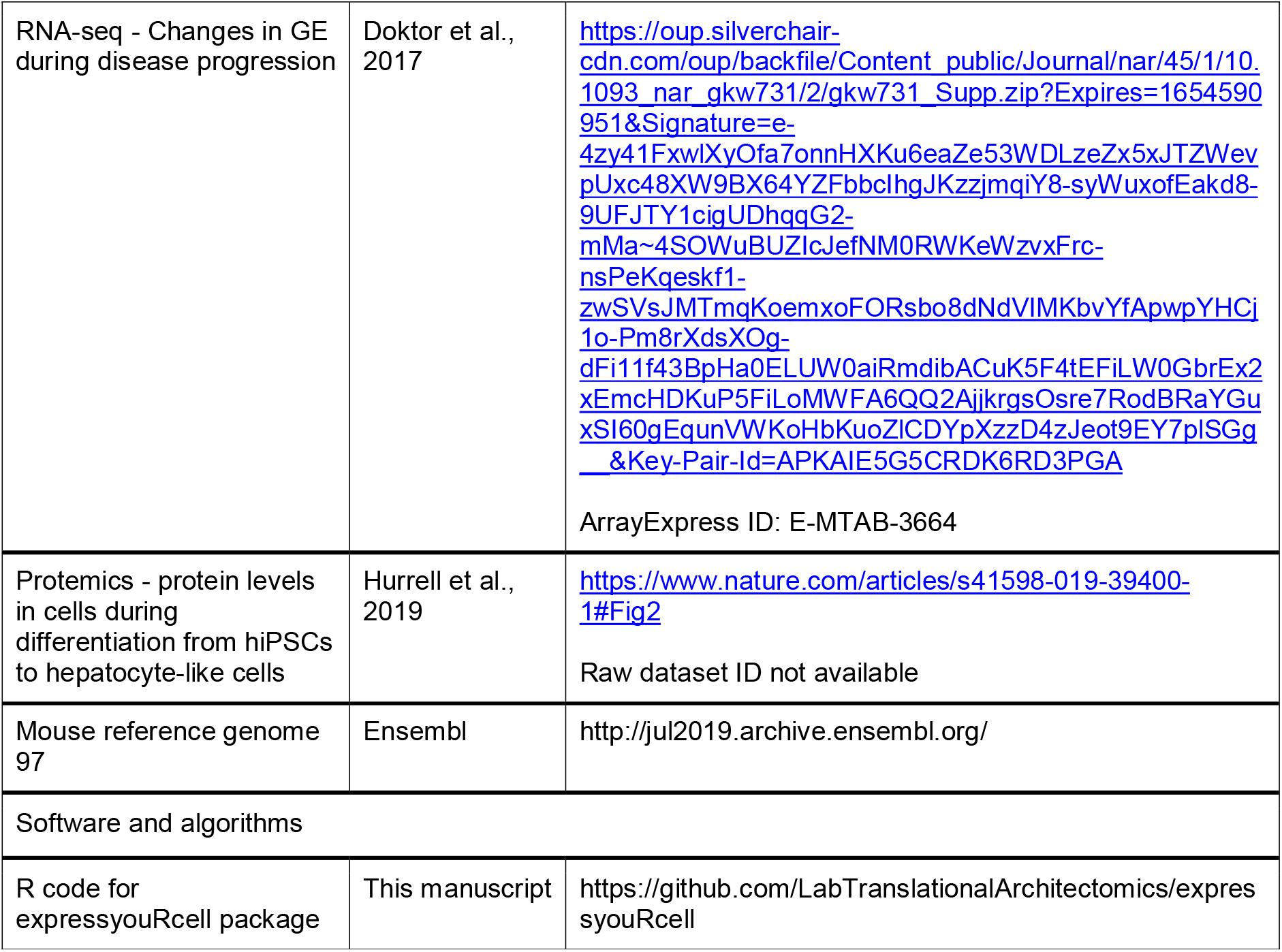

## ACKNOWLEDGMENT

The authors thank Ludovica Maria Ferrari for the graphical support, Emma Busarello and Christian Ramirez Amarilla for the comments.

## FUNDING

This work was supported by Telethon (reference no. GGP19115).

## AUTHOR CONTRIBUTIONS

M.P. developed the R package, organised all figures and wrote the draft of the paper. T.T. and F.L. advised on data interpretation and R package development. G.V. designed the work and obtained the funding. All authors wrote the paper and critically reviewed the manuscript.

## CONFLICT OF INTEREST

GV is scientific advisor of IMMAGINA Biotechnology s.r.l.

## Notes

https://www.frontiersin.org/articles/file/downloadfile/244243_supplementary-materials_datasheets_1_xlsx/octet-stream/Data%20Sheet%201.XLSX/1/244243

https://www.refine.bio/experiments/SRP055008/developing-mouse-cortex-rna-seq

https://static-content.springer.com/esm/art%3A10.1186%2Fs13024-018-0296-y/MediaObjects/13024_2018_296_MOESM2_ESM.xlsx

https://www.cell.com/cms/10.1016/j.neuron.2017.09.008/attachment/229d65b5-db13-442e-9721-fe2763746e2f/mmc7.xlsx

https://oup.silverchair-cdn.com/oup/backfile/Content_public/Journal/nar/45/1/10.1093_nar_gkw731/2/gkw731_Supp.zip?Expires=1654590951&Signature=e-4zy41FxwlXyOfa7onnHXKu6eaZe53WDLzeZx5xJTZWevpUxc48XW9BX64YZFbbcIhgJKzzjmqiY8-syWuxofEakd8-9UFJTY1cigUDhqqG2-mMa~4SOWuBUZIcJefNM0RWKeWzvxFrc-nsPeKqeskf1-zwSVsJMTmqKoemxoFORsbo8dNdVIMKbvYfApwpYHCj1o-Pm8rXdsXOg-dFi11f43BpHa0ELUW0aiRmdibACuK5F4tEFiLW0GbrEx2xEmcHDKuP5FiLoMWFA6QQ2AjjkrgsOsre7RodBRaYGuxSI60gEqunVWKoHbKuoZlCDYpXzzD4zJeot9EY7plSGg__&Key-Pair-Id=APKAIE5G5CRDK6RD3PGA

https://www.nature.com/articles/s41598-019-39400-1#Fig2

