## supplementary data for "Filming space-time changes of gene expression with *expressyouRcell*"

***Additional case studies - displaying longitudinal differential analyses***

*Additional case studies 1*

We took advantage of another RNA-seq dataset obtained by Wang and colleagues (7). This dataset is characterised by an experimental design with two covariates: time and disease condition. The authors performed a transcriptomic longitudinal differential analysis of microglial cells during a time span of 8 months. Microglial cells were isolated from control and rTg4510 mice, a murine model widely used for studying tauopathies such as Fronto-Temporal-Dementia (FTD) and Alzheimer’s disease. Differential analysis was performed between the two conditions at each time point. The authors took advantage of volcano plots to display the quantity and magnitude of differentially expressed genes (DEGs). They observed that within the first two time points, both the number of DEGs, and the magnitude of their fold changes are smaller compared with the results of differential analysis at 6 and 8 months. Using *expressyouRcell* we obtained 8 pictographs showing similar results. The pictographs show robust changes from month 4 onwards (**Figure** **S1A and movies S6-S7**).

Gene set enrichment analysis revealed an enrichment in the cellular membrane for down-regulated genes. In addition to this, ribosomes are also strongly enriched, in particular for up-regulated genes (**Figure** **S1B)**. Coherently, our pictographs generated with the option “gene set enrichment analysis” highlight these very same cellular compartments, confirming the presence of a strong enrichment for these terms, but also for cytoplasm and various other cellular localizations (**Figure** **S1C and movies S8-S9**).

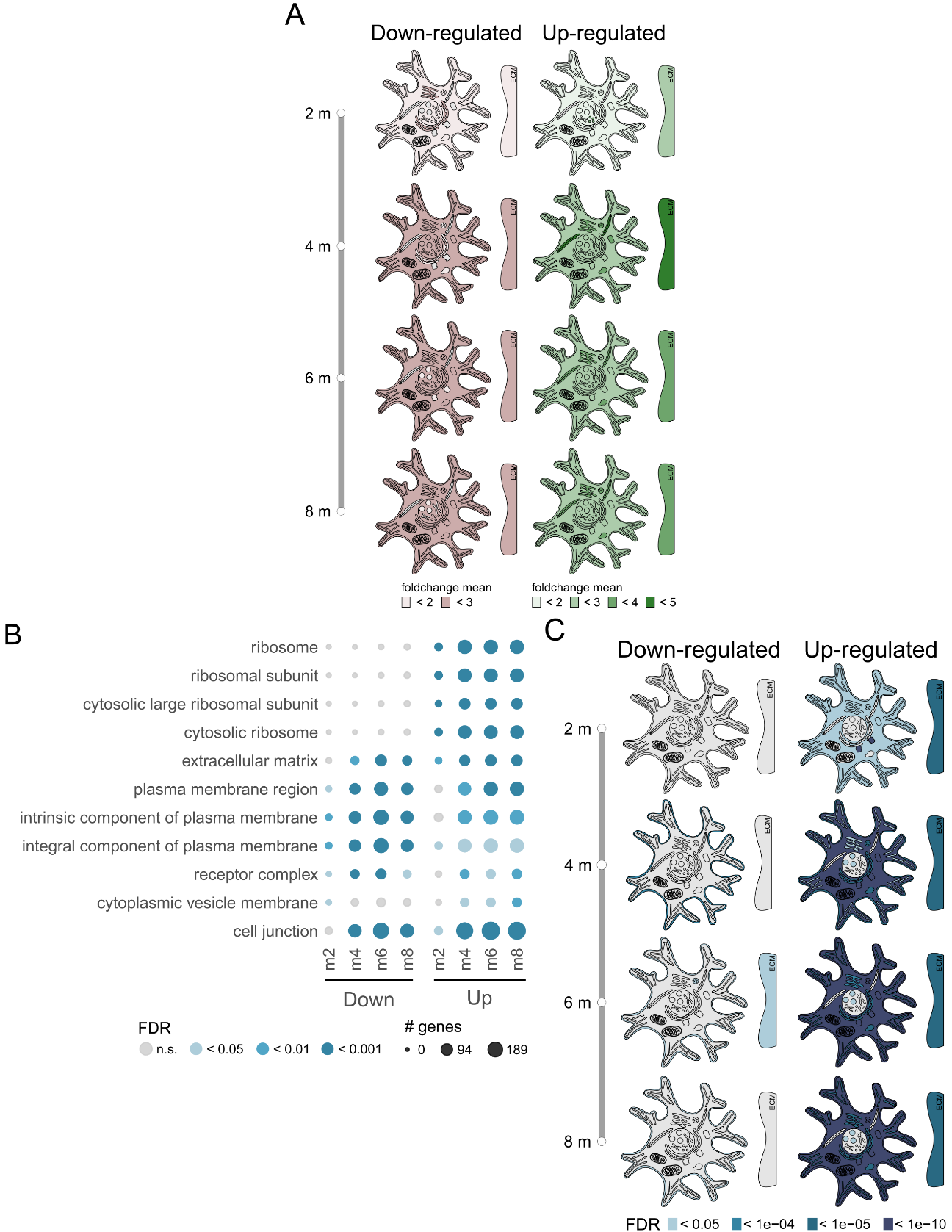

**Figure S1**. Related to STAR Methods. Comparisons between different visualisation methods on data from RNA-seq of microglial cells from (Wang et al. 2018). (A) Cell pictographs from RNA-seq of microglial cells. Colour shades of cellular compartments and organelles are based on logFC values from differential analysis. Up- and down-regulated genes were defined with the threshold described by (Wang et al. 2018) (i.e. abs(log2FC) > 1.5 and p-value < 0.05). (B) Enrichment analysis of Gene Ontology (Cellular Component) terms at indicated time point. Colours of dots indicate the significance of the enrichments. Dot sizes define the number of genes associated with each category.

(C) Cell pictographs from RNA-seq of microglial cells. Colour shades of cellular compartments and organelles are based on the significance of the enrichments. Up- and down-regulated genes were defined with the threshold described by (Wang et al. 2018) (i.e. abs(log2FC) > 1.5 and p-value < 0.05).

*Additional case studies 2*

We then applied *expressyouRcell* on a dataset based on an RNA-seq time-course analysis with six time points. Clements and collaborators (9) characterised the transcriptome of murine Schwann cells purified from intact nerves (iSC) and distal regions of nerves (dSC) following transection and investigated the transcriptional changes between the two conditions through differential gene expression analysis.

We used volcano plots to display the quantity and magnitude of differentially expressed genes (DEGs) identified between Schwann cells isolated from the intact and distal regions at each time point. We observed that within the first two time points, both the number of DEGs, and the magnitude of their fold changes, are smaller compared with the results of differential analysis starting from day 4. In general, we can also note that the number and magnitude of distal-enriched genes (**Figure S2A**, green dots) is higher with respect to intact-enriched genes (**Figure S2A**, red dots) at each time point.

We then used *expressyouRcell* to display outcomes of differential gene expression analysis (fold changes) through cellular pictograms (**Figure S2C**). For each time point, we obtained two cellular pictograms for representing fold changes of intact- (**Figure S2C**, red shaded pictograms) and distal-enriched genes (**Figure S2C**, green shaded pictograms), respectively. Compared with intact-enriched pictograms, we can observe that distal-enriched pictograms show a magnitude of distal-enriched genes larger than those enriched in Schwann cells from the intact region, coherently with volcano plot representations. With this visualisation we can appreciate the spatial localisation of gene expression changes and observe that, in particular at day 2, genes enriched in distal SC encode for proteins localised to the Golgi apparatus. On the other hand, genes with higher fold changes for intact SC are mainly associated with lysosomes at day 10.

Respectively on distal- and intact-enriched genes, we then performed gene set enrichment analysis restricted to the cellular components ontology, and reported the results of this analysis for each time point through heatmap visualisation (**Figure S3B**). Genes enriched in the intact Schwann cells region are mainly associated with the membrane cell region, in particular starting from day 4. On the other hand, starting from day 2, genes enriched in the intact Schwann cells are associated with the extracellular matrix and vesicles themes. We then generated cellular pictograms reporting results of gene set enrichment analysis for each time point, on genes enriched in the intact and distal Schwann cells, respectively (**Figure S2D and movies S14-S15**). Consistently with the heatmap visualisation, the enrichment for vesicles terms and for the extracellular matrix region clearly emerges across all the time points of distal Schwann cells associated genes.

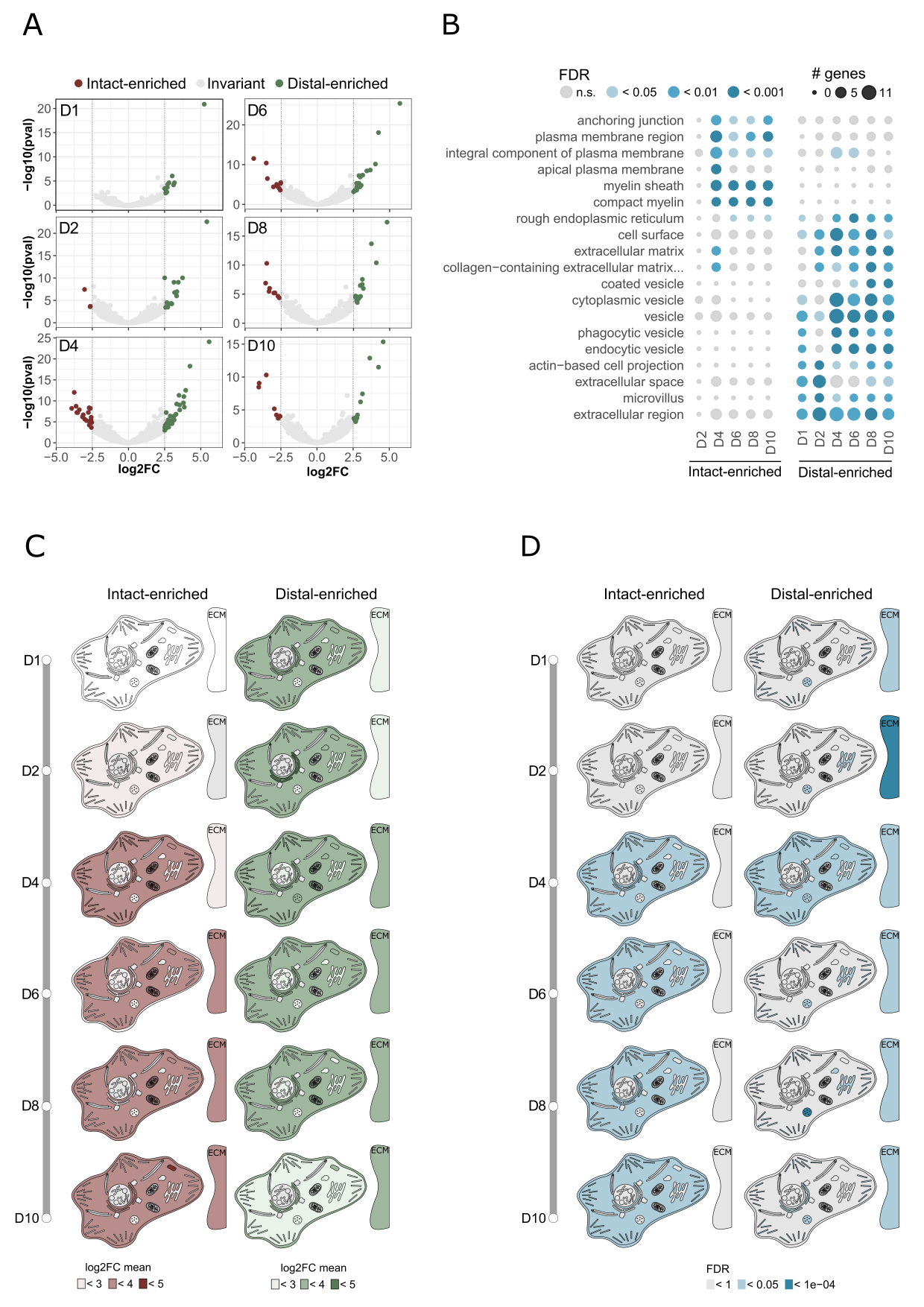

**Figure S2**. Related to STAR Methods. Comparisons between different visualisation methods on data from RNA-seq of Schwann cells from (Clements et al. 2017).

(A) Volcano plot of DEGs in distal Schwann cells relative to intact Schwann cells at indicated time point. Fold changes are plotted against the –log(p value). The vertical dashed lines indicate the 2.5 fold change cutoff thresholds.

(B) Enrichment analysis of Gene Ontology (Cellular Component) terms in all at indicated time points. The heatmap is coloured according to the significance of the enrichments and the number of genes associated to each category is reported.

(C) Cell pictographs from RNA-seq of Schwann cells. Colour shades of cellular compartments and organelles are based on logFC values from differential analysis. Up- and down-regulated genes were defined with the threshold described by (Clements et al. 2017). (i.e. abs(log2FC) > 2.5 and p-value < 0.05).

(D) Cell pictographs from RNA-seq of Schwann cells. Colour shades of cellular compartments and organelles are based on the significance of the enrichments. Up- and down-regulated genes were defined with the threshold described by (Clements et al. 2017) (i.e. abs(log2FC) > 2.5 and p-value < 0.05).

*Additional case studies 3*

We applied *expressyouRcell* on RNA-seq datasets of a complex experimental design with three covariates: condition, time and tissue (8). The authors profiled gene expression in different tissues (brain, spinal cord, liver and muscle) and two stages (pre- and early-symptomatic) of disease in a mouse model of Spinal Muscular Atrophy (8). Differential analysis showed few changes at the pre-symptomatic stage, while at the early-symptomatic stage a larger number of differentially expressed genes was found considering multiple tissues and different conditions. Consistently with Doktor and collaborators (8), a stronger enrichment for differentially expressed genes at the early-symptomatic with respect to the pre-symptomatic stage can be visualised by *expressyouRcell* in all tissues (**Figure S3 and movies S10-S13**).

The authors observed a significant enrichment of down-regulated genes in the cell cycle KEGG pathway in the spinal cord and muscle at the early-symptomatic stage. Interestingly, our visualisation highlights also an additional and strong enrichment for chromosomes. Although the authors did not observe significantly enriched terms for up-regulated genes, cellular pictographs show enrichments in several cellular compartments and organelles, particularly at an early-symptomatic stage of disease. The differences between our representation and their results might be explained by the use of diverse databases to obtain the gene set enrichment analysis (i.e. the cellular components ontology in our case, and KEGG in their work).

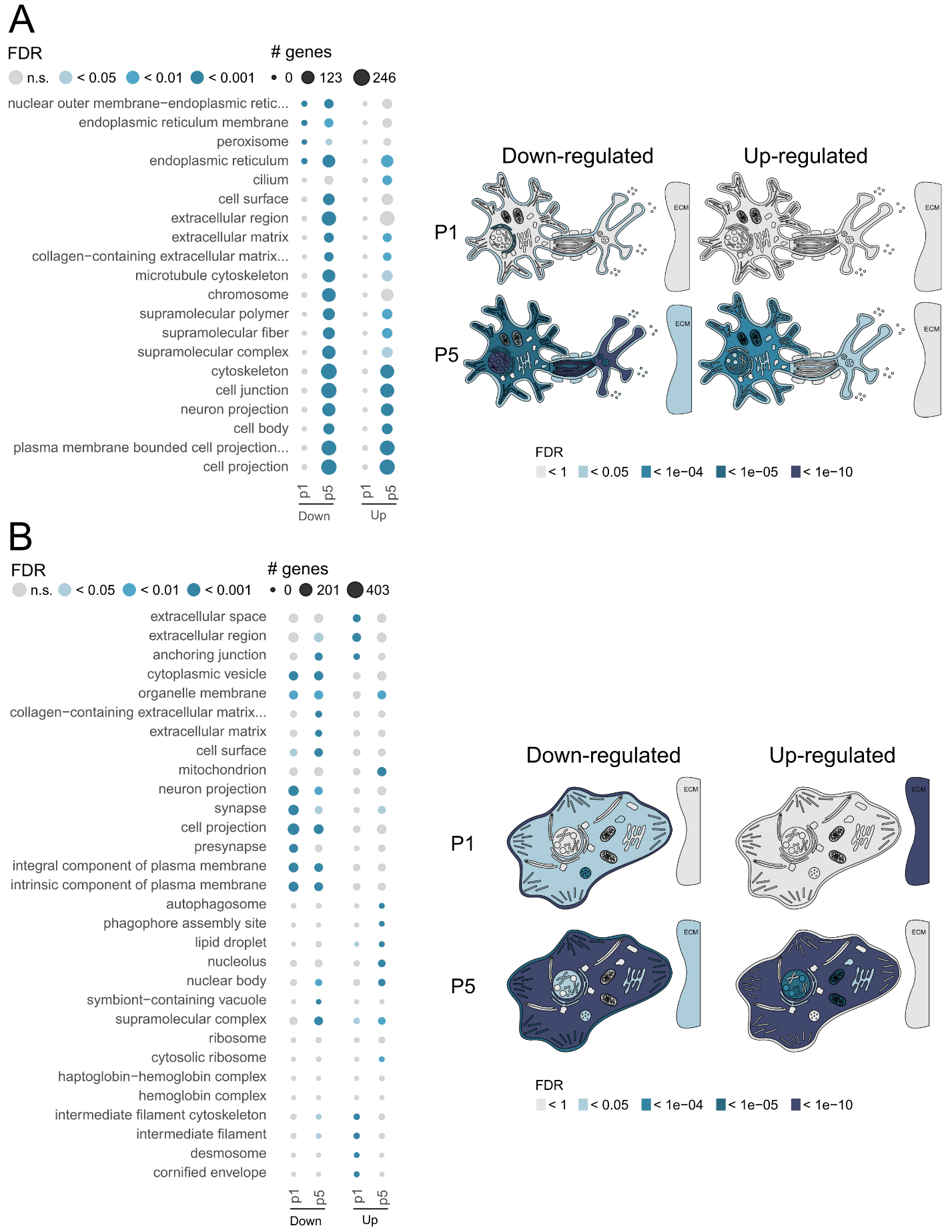

**Figure S3**: Related to STAR Methods. Comparisons between different visualisation methods on data from RNA-seq on spinal cord (A) and muscle tissue (B) from (Doktor et al. 2017).

(A) Left. Enrichment analysis of Gene Ontology (Cellular Component) terms at indicated time point for differentially expressed genes identified in the spinal cord. The heatmap is coloured according to the significance of the enrichments and the number of genes associated to each category is reported. Right. Neuron pictographs from RNA-seq on the spinal cord. Colour shades of cellular compartments and organelles are based on the significance of the enrichments. Up- and down-regulated genes were defined with the threshold described by (Doktor et al. 2017) (i.e. abs(log2FC) > 0.12 and FDR < 0.1).

(B) Left. Enrichment analysis of Gene Ontology (Cellular Component) terms at indicated time point for differentially expressed genes identified in muscle. The heatmap is coloured according to the significance of the enrichments and the number of genes associated to each category is reported. Right. Cell pictographs from RNA-seq on muscle. Colour shades of cellular compartments and organelles are based on the significance of the enrichments. Up- and down-regulated genes were defined with the threshold described by (Doktor et al. 2017) (i.e. abs(log2FC) > 0.12 and FDR < 0.1).

*Additional case studies 4*

To display the versatility of our tool, we applied expressyouRcell to a proteomics dataset (Hurrell et al. 2019). Here the authors analysed proteins during differentiation of human induced pluripotent stem cells (hiPSCs) into hepatocytes-like cells (HLCs) at eight time points. For each time point we generated pictographs of generic mammalian cell (**Figure S4**) based on protein levels. We can observe that the protein levels signal increase and decrease in waves across time, across various cellular localizations. Starting from the first day, high protein abundances are generally spread across the entire cell, while at days 3, 7 and 25 higher protein levels are mainly focused within specific organelles. We then applied the animate function to generate dynamic cellular pictographs, which clearly support the visualization of protein levels movement across time point and cellular compartments (**Movie S16**).

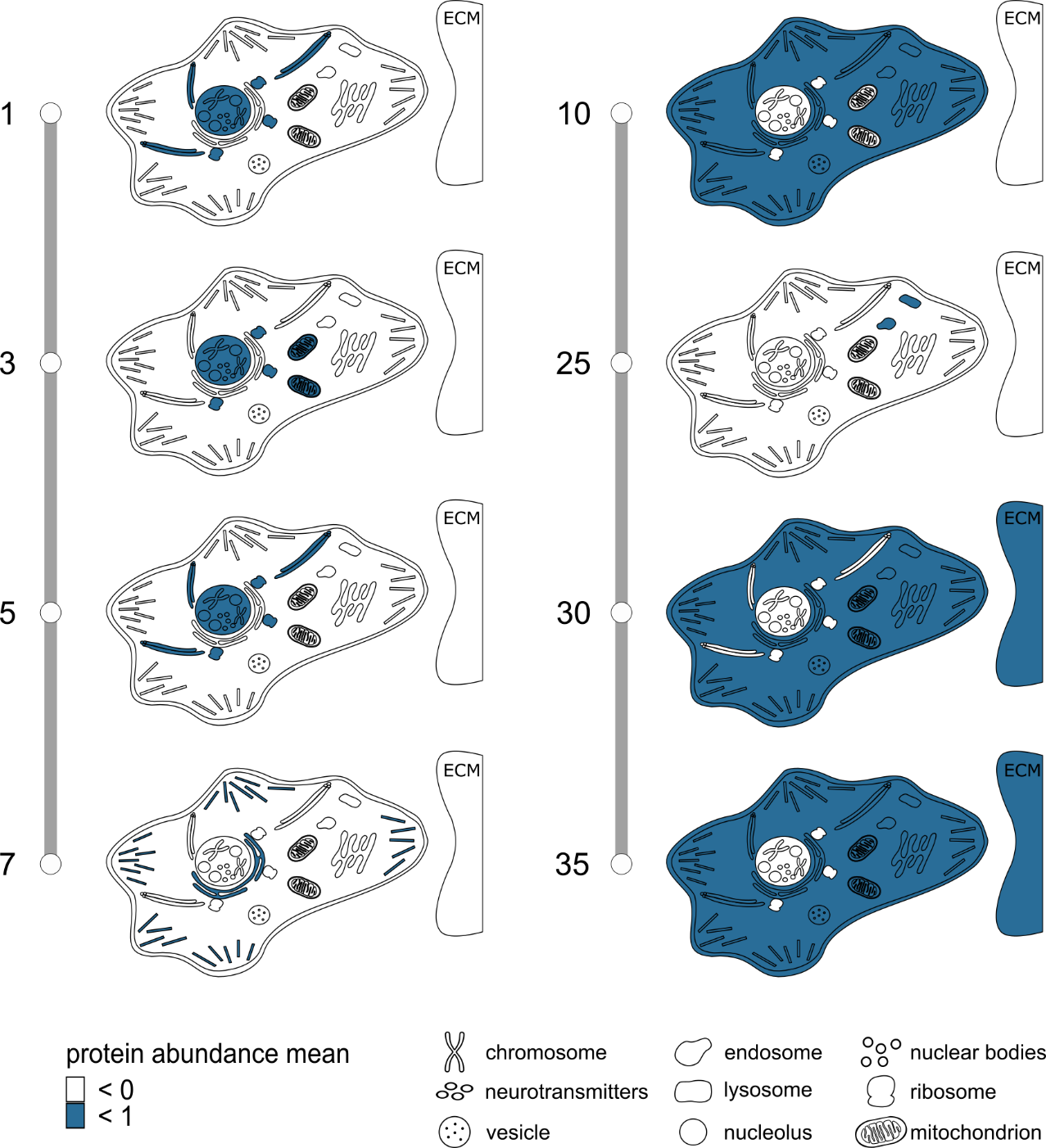

**Figure S4**: Related to STAR Methods. Mammalian cell pictographs from proteomic of hiPSCs. Colour shades of cellular compartments and organelles are based on protein abundance levels.

***Quantification and statistical analysis***

Colour shades of pictographs based on the significance of the enrichments were defined by p-values from one side Fisher’s test.

| Cell structures | Cell types |
| --- | --- |
| Membrane, nucleoplasm, endoplasmic  reticulum, Golgi apparatus, microtubules, actin,  mitochondrion, vesicle, nucleolus, nuclear body,  lysosome, endosome, chromosome,  extracellular region, ribosome | Neuron, fibroblast, generic mammalian cell,  microglia |
| Synapse, axon, cell body, synaptic membrane,  myelin sheath | Neuron |

**Table S1.** List of organelles and cellular compartments included in the cellular pictographs.

**Movie S1.** Animated neuron pictograph from TRAP-seq of cortical neurons - down-regulated genes. The movie includes all the three time points, from 30’ to 120’. Colour shades of cellular compartments and organelles are based on logFC values of down-regulated genes from differential analysis. Up- and down-regulated genes were defined with the threshold described by (6) (i.e. abs(logFC) > 0.4 and FDR < 0.1).

**Movie S2**. Animated neuron pictograph from TRAP-seq of cortical neurons - up-regulated genes. The movie includes all the three time points, from 30’ to 120’. Colour shades of cellular compartments and organelles are based on logFC values of up-regulated genes from differential analysis. Up- and down-regulated genes were defined with the threshold described by (6) (i.e. abs(logFC) > 0.4 and FDR < 0.1).

**Movie S3.** Animated neuron pictograph from TRAP-seq of cortical neurons - down-regulated genes. The movie includes all the three time points, from 30’ to 120’. Colour shades of cellular compartments and organelles are based on the significance of the enrichments of down-regulated genes. Up- and down-regulated genes were defined with the threshold described by (6) (i.e. abs(logFC) > 0.4 and FDR < 0.1).

**Movie S4**. Animated neuron pictograph from TRAP-seq of cortical neurons - up-regulated genes. The movie includes all the three time points, from 30’ to 120’. Colour shades of cellular compartments and organelles are based on the significance of the enrichments of up-regulated genes. Up- and down-regulated genes were defined with the threshold described by (6) (i.e. abs(logFC) > 0.4 and FDR < 0.1).

**Movie S5.** Animated neuron pictograph from RNA-seq of cortical neurons. The movie is created from the E14.5 stage to the 21st month. Colour shades of cellular compartments and organelles are based on gene expression levels (measured in TPM).

**Movie S6**. Animated microglia pictograph from RNA-seq of microglial cells - down-regulated genes. The movie includes all the four time points, from 2 months to 8 months. Colour shades of cellular compartments and organelles are based on logFC values of down-regulated genes from differential analysis. Up- and down-regulated genes were defined with the threshold described by (7) (i.e. abs(log2FC) > 1.5 and p-value < 0.05).

**Movie S7**. Animated microglia pictograph from RNA-seq of microglial cells - up-regulated genes. The movie is created including all the four time points, from 2 months to 8 months. Colour shades of cellular compartments and organelles are based on logFC values of up-regulated genes from differential analysis. Up- and down-regulated genes were defined with the threshold described by (7) (i.e. abs(log2FC) > 1.5 and p-value < 0.05).

**Movie S8**. Animated microglia pictograph from RNA-seq of microglial cells - down-regulated genes. The movie includes all the four time points, from 2 months to 8 months. Colour shades of cellular compartments and organelles are based on the significance of the enrichments of down-regulated genes. Up- and down-regulated genes were defined with the threshold described by (7) (i.e. abs(log2FC) > 1.5 and p-value < 0.05).

**Movie S9**. Animated microglia pictograph from RNA-seq of microglial cells - up-regulated genes. The movie includes all the four time points, from 2 months to 8 months. Colour shades of cellular compartments and organelles are based on the significance of the enrichments of up-regulated genes. Up- and down-regulated genes were defined with the threshold described by (7) (i.e. abs(log2FC) > 1.5 and p-value < 0.05).

**Movie S10.** Animated cell pictographs from RNA-seq of Schwann cells - down-regulated genes. The movie includes six time points, from D1 to D10. Colour shades of cellular compartments and organelles are based on the significance of the enrichments of down-regulated genes. Up- and down-regulated genes were defined with the threshold described by (9) (i.e. abs(log2FC) > 2.5 and p-value < 0.05).

**Movie S11.** Animated cell pictographs from RNA-seq of Schwann cells - up-regulated genes. The movie includes six time points, from D1 to D10. Colour shades of cellular compartments and organelles are based on the significance of the enrichments of up-regulated genes. Up- and down-regulated genes were defined with the threshold described by (9) (i.e. abs(log2FC) > 2.5 and p-value < 0.05).

**Movie S12.** Animated neuron pictograph from RNA-seq of spinal cord tissues - down-regulated genes. The movie includes two time points, from PND1 to PND5. Colour shades of cellular compartments and organelles are based on the significance of the enrichments of down-regulated genes. Up- and down-regulated genes were defined with the threshold described by (8) (i.e. abs(log2FC) > 0.12 and FDR < 0.1).

**Movie S13**. Animated neuron pictograph from RNA-seq of spinal cord tissues - up-regulated genes. The movie includes two time points, from PND1 to PND5. Colour shades of cellular compartments and organelles are based on the significance of the enrichments of up-regulated genes. Up- and down-regulated genes were defined with the threshold described by (8) (i.e. abs(log2FC) > 0.12 and FDR < 0.1).

**Movie S14**. Animated cell pictograph from RNA-seq of muscle tissue - down-regulated genes. The movie includes two time points, from PND1 to PND5. Colour shades of cellular compartments and organelles are based on the significance of the enrichments of down-regulated genes. Up- and down-regulated genes were defined with the threshold described by (8) (i.e. abs(log2FC) > 0.12 and FDR < 0.1).

**Movie S15.** Animated cell pictograph from RNA-seq of muscle tissue - up-regulated genes. The movie includes two time points, from PND1 to PND5. Colour shades of cellular compartments and organelles are based on the significance of the enrichments of up-regulated genes. Up- and down-regulated genes were defined with the threshold described by (8) (i.e. abs(log2FC) > 0.12 and FDR < 0.1).

**Movie S16**. Animated cell pictograph from protein abundance levels of hiPSCs. The movie is created from day 1 to 35. Colour shades of cellular compartments and organelles are based protein abundance levels.
